# Analyzing contentious relationships and outlier genes in phylogenomics

**DOI:** 10.1101/115774

**Authors:** Joseph F. Walker, Joseph W. Brown, Stephen A. Smith

## Abstract

Recent studies have demonstrated that conflict is common among gene trees in phylogenomic studies, and that less than one percent of genes may ultimately drive species tree inference in supermatrix analyses. Here, we examined two datasets where supermatrix and coalescent-based species trees conflict. We identified two highly influential “outlier” genes in each dataset. When removed from each dataset, the inferred supermatrix trees matched the topologies obtained from coalescent analyses. We also demonstrate that, while the outlier genes in the vertebrate dataset have been shown in a previous study to be the result of errors in orthology detection, the outlier genes from a plant dataset did not exhibit any obvious systematic error and therefore may be the result of some biological process yet to be determined. While topological comparisons among a small set of alternate topologies can be helpful in discovering outlier genes, they can be limited in several ways, such as assuming all genes share the same topology. Coalescent species tree methods relax this assumption but do not explicitly facilitate the examination of specific edges. Coalescent methods often also assume that conflict is the result of incomplete lineage sorting (ILS). Here we explored a framework that allows for quickly examining alternative edges and support for large phylogenomic datasets that does not assume a single topology for all genes. For both datasets, these analyses provided detailed results confirming the support for coalescent-based topologies. This framework suggests that we can improve our understanding of the underlying signal in phylogenomic datasets by asking more targeted edge-based questions.

## INTRODUCTION

Recent phylogenomic studies have shown that small changes to a dataset or the methods used to analyze a dataset can yield conflicting hypotheses at particular recalcitrant relationships with high support (i.e., 100% support from nonparametric bootstrap (BS) or posterior probability (PP) values). Prominent examples of this include many charismatic lineages such as the root of placental mammals (Morgan et al. 2013; Romiguier et al. 2013), early branching within Neoaves (Jarvis et al. 2014; Prum et al. 2015), and the earliest diverging lineage of extant angiosperms (Zanis et al. 2002; Wickett et al. 2014; Xi et al. 2014). The resolution of these relationships is critical to understanding the evolutionary history of their respective clades (e.g., patterns of biochemical, morphological, and life history evolution).

Finding the underlying causes of uncertainty in phylogenetic datasets is an essential step toward resolving problematic relationships. Recently, authors have developed means of exploring conflict between gene trees and species trees specifically for phylogenomic datasets (Salichos et al. 2014; Smith et al. 2015; Kobert et al. 2016), aiding in the identification of regions of species trees with considerable uncertainty despite strong statistical support from traditional support measures. Two studies have shown that the disproportionate influence of just one or two “outlier genes” on a supermatrix analysis is capable of driving tree topology inference (Brown and Thomson 2017; Shen et al. 2017). Using a Bayes factor approach Brown and Thomson (2017) reanalyzed a series of published datasets and found that the transcriptome data from Chiari et al. (2012) contained outlier genes. When outlier genes were included in phylogenetic reconstruction, a clade of turtles+crocodilians was inferred to be sister to birds with 100% PP. The same topology was previously inferred using ML with nucleotide data in the original study by Chiari et al. (2012) but was dismissed in favor of a coalescent reconstruction that placed turtles sister to birds+crocodilians. When Brown and Thomson (2017) removed the outlier genes, the reduced supermatrix inferred the same topology as the coalescent reconstruction with 100% PP. Another recently published study compared gene-wise likelihoods across multiple topologies to examine contentious relationships across the tree of life and found disproportionate influence of genes at all contentious relationships examined (Shen et al. 2017).

While such studies have highlighted several issues concerning phylogenomic conflict within datasets, these are early steps and several of these approaches have limitations that may limit our ability to identify phylogenetic support for particular relationships. For example, some of these analyses may incur significant runtimes that may limit more extensive dataset exploration or be a barrier for larger datasets. Also, these analyses are often performed on a small number (e.g., ∼2) of alternative topologies (e.g., Castoe et al. 2009; Smith et al. 2011; Shen et al. 2017), and like typical supermatrix analyses, most explicitly assume that all genes share a topology. However, given widespread gene tree discordance (e.g., due to incomplete lineage sorting [ILS] and other processes), it may be more realistic to assume that many alternative topologies are supported within datasets (e.g., Smith et al. 2015; Pease et al. 2016; Walker et al. 2017). Coalescent species tree methods relax this assumption but typically assume that gene tree discordance is the result of ILS (but see Boussau et al. 2013). The computational burden of large datasets also typically limits these coalescent analyses to Maximum Quartet Support Species Tree (MQSST) methods (Mirarab and Warnow 2015) that have additional simplifying assumptions.

If the research question involves a small number of relationship and not the entirety of the tree, it may be more appropriate to examine targeted edges instead of resolved topologies (Lee and Hugall 2003). Here, we describe a fast analysis framework, maximum gene-wise edge (MGWE) analysis. This framework facilitates the examination of contentious edges in phylogenomic datasets without the requirement that each gene share the same topological resolution. We compare results from two-topology gene-wise log-likelihood and MGWE analyses for vertebrate (Chiari et al. 2012; Brown and Thomson 2017) and carnivorous Caryophyllales datasets (Walker et al. 2017) (hereafter referred to as the carnivory dataset). Both datasets contain contentious relationships, outlier genes, and, in their respective original studies, the authors dismissed the supermatrix topology for the topology inferred using a coalescent method. In both cases, we find that the use of an edge based approach results in stronger support for the topology hypothesized to be correct by researchers in the original study.

## METHODS

### Data collection

We obtained the 248 genes that were codon-aligned and analyzed by Brown and Thomson (2017) from the Dryad deposit (http://dx.doi.org/10.5061/dryad.8gm85) of the original study (Chiari et al. 2012) that focused on resolving the placement of turtles among amniotes. The coding DNA sequences of the 1237 one-to-one orthologs from Walker et al. (2017) to infer the relationships among carnivorous Caryophyllales (Eudicots: Superasterids) are available from Dryad (http://datadryad.org/resource/doi:10.5061/dryad.vn730). All programs used in this analysis may be found at: https://github.com/jfwalker/MGWE.

### Species trees

Brown and Thomson (2017) used Bayesian analyses to obtain the topologies from the Chiari et al. (2012) data set. As our study focused on the use of maximum likelihood (ML) for detecting overly influential genes, we ensured that ML phylogenetic reconstruction would recapitulate the previous species tree results. To construct a supermatrix tree for the vertebrate dataset, the 248 individual vertebrate genes used in Brown and Thomson (2017) were concatenated using the Phyx program pxcat (Brown et al. 2017). The species tree was inferred in RAxML v8.2.10 (Stamatakis 2014) using the GTR+ Γ model of evolution, and edge support was assessed from 200 rapid bootstrap replicates. Supermatrix trees for the vertebrate dataset were inferred both with all genes present, and again with the previously identified two outlier genes (8916 and 11434) removed (see below). The ML tree inferred from all the data from the carnivory dataset was downloaded from (http://dx.doi.org/10.5061/dryad.vn730) while a novel ML tree was inferred from a reduced supermatrix that excluded two highly influential genes (cluster575 and cluster3300; see below).

### Gene tree construction and analysis of conflict

Individual gene trees for both datasets were inferred using ML with the GTR+ Γ model of evolution as implemented in RAxML. SH-like analyses (Anisimova et al. 2011), as implemented in RAxML, were performed to assess gene tree edge support. As this analysis examines alternative topologies by nearest-neighbor interchange (NNI), it is possible that during the analysis a topology with a higher likelihood is found (i.e., an ‘NNI-optimal’ topology). When a better topology was found, that topology was used in downstream analyses. We used the pxrr program in the Phyx package (Brown et al. 2017) to root all gene trees on the outgroup (*Protopterus* for the vertebrate dataset, and *Beta vulgaris* and *Spinacia oleraceae* for the carnivory dataset) and we excluded gene trees where an outgroup was not present. We mapped conflict onto the supermatrix tree using phyparts (Smith et al. 2015) with SH-like support of < 80 treated as uninformative. We chose 80 as a support cutoff as 95 has been shown to be overly conservative (Guindon et al. 2010). Gene tree conflict was visualized using the script phypartspiecharts.py (available from https://github.com/mossmatters/MJPythonNotebooks). We conducted more detailed conflict analyses used for edge comparisons discussed below using pxbp as part of the Phyx package (Brown et al. 2017).

### Calculating two-topology gene-wise log-likelihoods

The alternate topologies (supermatrix and coalescent) and data matrices for the vertebrate and carnivory datasets were obtained from the original studies, Chiari et al. (2012) and Walker et al. (2017), respectively. We calculated site-wise log-likelihood scores for the two topologies in RAxML using the GTR+ Γ model of evolution, with the data partitioned by gene. The differences in site-wise log-likelihoods between the candidate topologies were then calculated using scripts available from https://github.com/jfwalker/MGWE.

### Maximum gene-wise edge calculations

In addition to pairwise topological comparisons, we also examined the maximum gene-wise edges (MGWE) (Fig 1.). For a single gene and a single focal edge, the MGWE is the resolution among a set of alternative resolutions for the focal edge that has the highest likelihood from among a set of topologies (more details can be found below). The set of topologies can be determined *a priori* or based on constrained phylogenetic analyses. With this approach, genes are not required to share the same topology even if genes have the same MGWE. This contrasts with a standard shared topology comparison where the topology for each gene would be required to be the same (e.g., supermatrix vs. coalescent topology). Therefore, the MGWE approach allows for genes to have conflicting relationships outside of the edge of interest whether or not they agree with the resolution for the edge of interest. Here, we compared the MGWE for sets of alternative and conflicting edges in order to determine if, by relaxing the requirement for each gene to share the topology, we gain insight into the signal for conflicting relationships.

**Figure 1.**
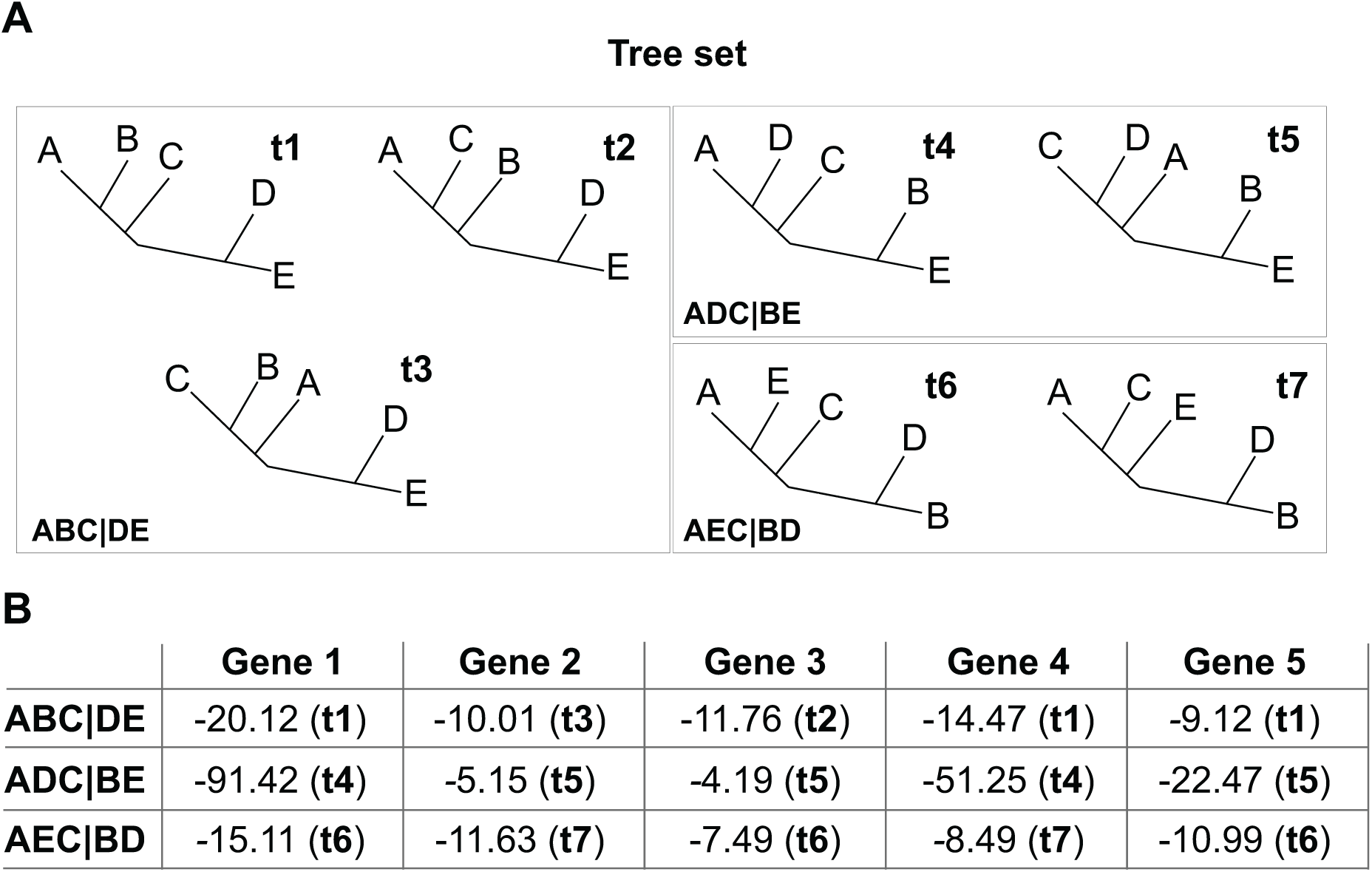
Outline for the MGWE procedure. (A) A tree set is depicted with trees numbered. Trees that are concordant for the edge of interest are grouped in boxes with each box representing a CADRE. The concordant edge of interest is denoted at the bottom left-hand corner of each box. (B) A table showing the highest likelihood for each edge calculated from the relevant CADRE and the tree (in parentheses) on which that likelihood was calculated. The MGWE would be the edge for each gene with the highest likelihood.

As mentioned above, the set of topologies that may be used to calculate MGWEs could be determined *a priori* or based on constrained phylogenetic reconstruction analyses. Here, we restricted the tree space under consideration by circumscribing a set of empirically supported topologies (TREESET) consisting of the supermatrix-inferred topology, coalescent inferred topology, and individual gene trees that contained all taxa. For each edge set (i.e., a particular edge and the dominant alternative edges) examined, we pooled trees that were concordant for a particular resolution involving the focal taxa in question for the edge set. Here, for simplicity, we call this set of trees that are concordant for a particular relationship a CADRE. Thus, there was a CADRE for each resolution for an edge of interest. We then calculated the maximum likelihood for each gene on each topology in the TREESET.

We calculated the MGWEs by retaining the likelihood for the topology with the highest likelihood for each CADRE across all the genes. This became the representative likelihood for that CADRE. The CADRE with the highest likelihood for the gene determined which resolution was the MGWE for that gene.

We then compared this more complex model, allowing for each gene to have a different topology and branch lengths, to the model assuming the shared supermatrix and coalescent topologies. To do this, we calculated the AIC and AICc scores for each CADRE as the summed likelihoods are not comparable given the differences in the number of parameters between the respective models (Theobald 2010; Posada and Buckley 2004). The parameters, *k*, were calculated based on the number of taxa in each gene, *n*, and the number of genes in the analysis, *g*. For a single gene, there were 2×*n*– 3 branch length parameters and 9 parameters for the GTR + Γ model of evolution (5 substitution parameters, 1 among-site rate heterogeneity parameter, and 3 estimated base frequencies parameters). The topology was not considered a parameter (Felsenstein, 1983; Yang et al. 1995), when calculating the AIC or AICc scores. The AICc score included a correction for the total number of sites in the supermatrix.

We compared the AIC and AICc scores of several alternative models. First, we ran a standard supermatrix ML analyses assuming a single set of branch lengths on one topology and model parameters unlinked across genes with a GTR + Γ model of evolution (2×*n* – 3 + 9 ×*g* parameters). We also conducted a supermatrix analysis allowing the branches to be unlinked across genes including 2 × *n* – 3 + 9 parameters for every partition and the total parameters being the sum of all parameters for each partition. For this analysis, the number of parameters were the same as those calculated for the CADRE analysis.

Here, we focused on addressing conflicting signal between edges of interest and so the increase in the number of parameters (i.e., a full set for each gene) was considered to be acceptable given our emphasis on gene trees comparisons. However, future work could attempt to limit the expansion of the number of parameters for each CADRE by sharing branch length estimates or model parameters across genes. The code for this analysis is available at https://github.com/jfwalker/MGWE.

### Testing for paralogy in carnivory dataset

The homolog trees created from amino acid data in the study by Walker et al. (2017) were downloaded from Dryad (http://datadryad.org/resource/doi:10.5061/dryad.vn730). We matched the sequences from the outlier genes to their corresponding sequence in the amino acid homolog trees. This allowed us to examine whether a nucleotide cluster contained homology errors that may be exposed by the slower evolving amino acid dataset.

## RESULTS

### Gene tree conflict and log-likelihood analysis reveals genes of disproportionate influence

Our ML analysis of the vertebrate dataset recovered the same supermatrix topology (Fig. 2) as found with ML by Chiari et al. (2012) and Bayesian inference by Brown and Thomson (2017). The difference in log-likelihood between the supermatrix and the coalescent-based Maximum Quartet Support Species Tree (hereafter referred to as coalescent) topologies for the vertebrate dataset was 4.01. Ninety-three of 248 gene trees could be rooted on the outgroup *Protopterus* and only five of these had all taxa represented (Supplementary Table 1). We found low support for relationships within gene trees (SH <80) and substantial gene tree conflict (Fig. 2). Of the gene trees with high support (SH >80), seven resolved turtles+crocodilians as sister to birds (hereafter referred to as the vertebrate supermatrix topology) and nine resolved crocodilians+birds sister to turtles (hereafter referred to as the vertebrate coalescent topology).

**Figure 2.**
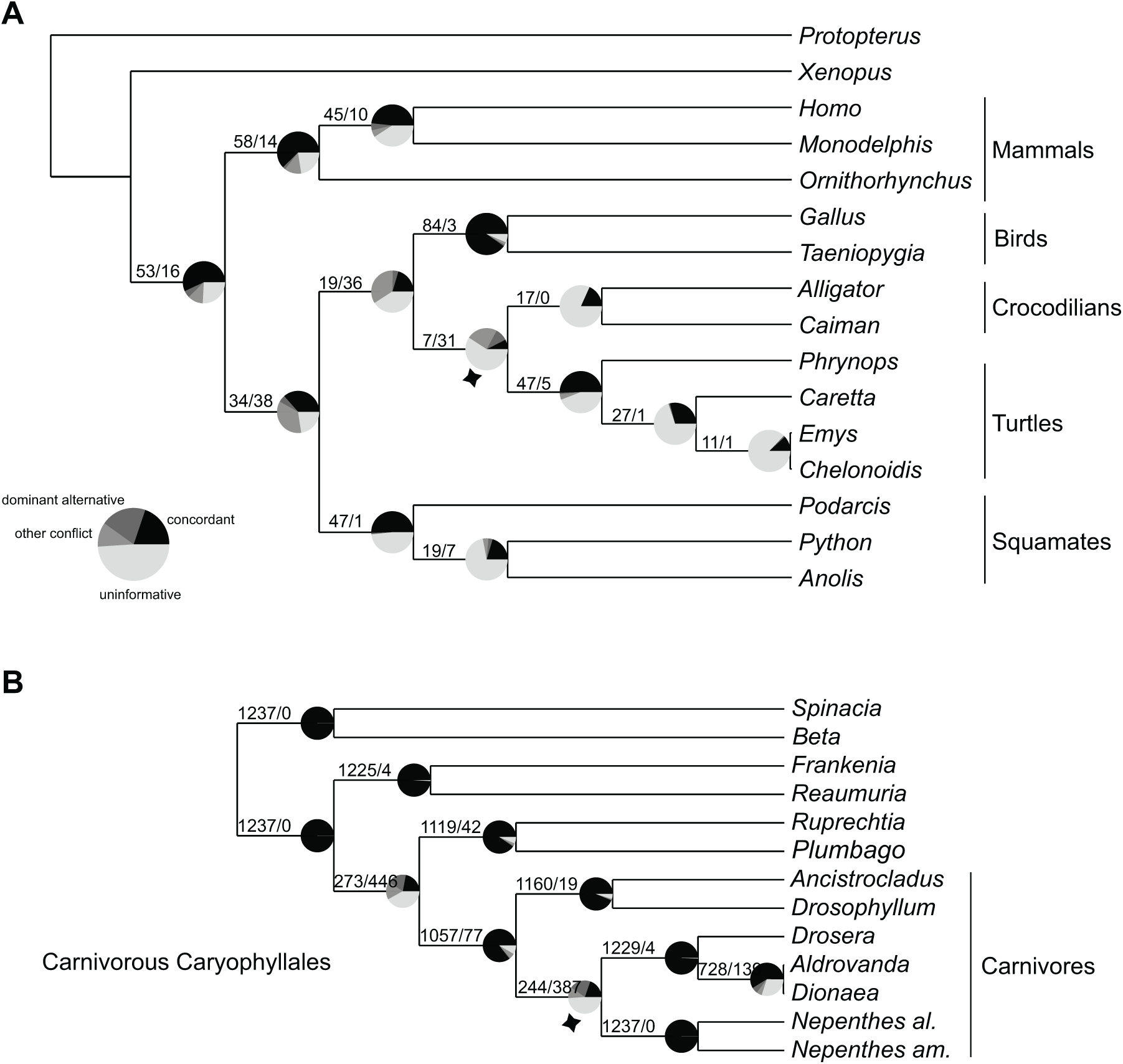
Maximum likelihood trees inferred by RAxML for the Chiari et al. 2012 (vertebrate) and Walker et al. 2017 (carnivorous Caryophyllales) datasets. Conflict analysis for the vertebrate (A) and carnivory (B) datasets. The vertebrate analysis includes the 93 genes that contained the outgroup (*Protopterus*), and the carnivory analysis includes 1237 genes all of which had the outgroups (*Spinacia oleraceae* and *Beta vulgaris*). Black represents gene trees that are concordant with the relationship, the lightest grey represents uninformative genes (SH-like < 80 or no taxon representation for the edge), dark grey represents the dominant alternate topology, and light grey represents all other conflict. Numbers on edges represent concordance/conflict. Bold numbers at the nodes of the vertebrate dataset correspond to edge numbers in Supplementary Table 1.

The two-topology gene-wise log-likelihood comparison showed that 105 genes had a higher likelihood score for the vertebrate supermatrix topology while 143 supported the vertebrate coalescent topology (Figs. 3A, 4A). Two genes (ENSGALG00000008916 and ENSGALG00000011434, referred to here as 8916 and 11434, respectively), appeared as outliers, exhibiting a disproportionate influence on the overall likelihood of the supermatrix (Fig. 3A). The outlier genes identified with maximum likelihood analyses matched those previously identified as outliers using Bayes factors (Brown and Thomson 2017). These two genes both supported the vertebrate supermatrix topology with log-likelihood scores of 79.55 and 46.01 greater than the alternative coalescent tree topology, respectively. The difference in log-likelihood between the two topologies of the non-outlier genes ranged from 0.006 to 19.891 with an average of 3.31 for all genes in the analysis. The removal of the vertebrate genes 8916 and 11434, as shown by Brown and Thomson (2017), recovered the coalescent topology, albeit with low bootstrap support (BS = 12; Supplementary Fig. 1).

**Figure 3.**
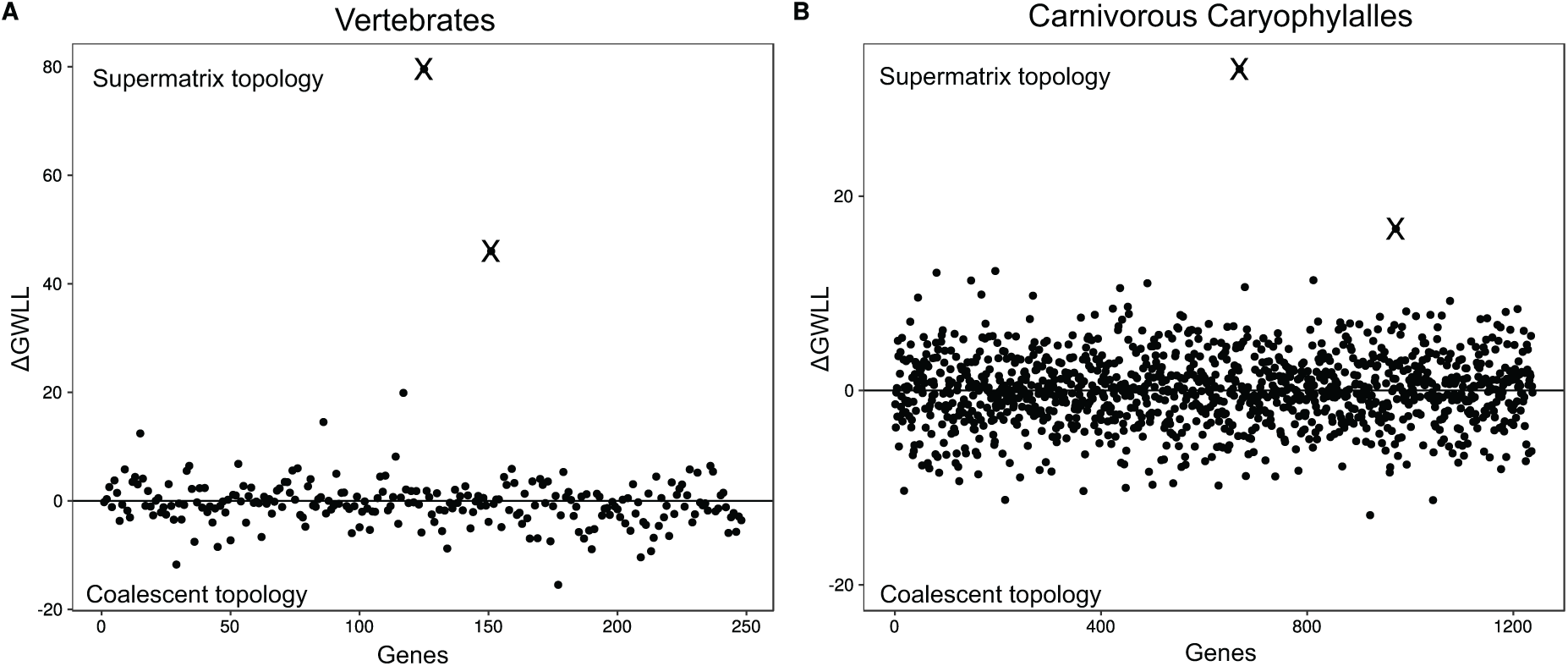
Identification of outlier genes using gene-wise likelihood comparison. A&B) Show the results of the two-topology gene-wise log-likelihood (GWLL) comparison on the vertebrate and carnivory dataset, respectively, using the coalescent (negative values) and supermatrix (positive values) topologies as the comparison. The genes identified as outliers from the analysis are marked with an X.

Previous work on the carnivory dataset demonstrated that the placement of the *Ancistrocladus+Drosophyllum* clade (Fig. 2) contained significant conflict and was strongly influenced by species sampling (Walker et al. 2017). The log-likelihood difference between the supermatrix and coalescent topologies was 74.94 in favor of the former. The two-topology log-likelihood comparison between the dominant topologies on the carnivory dataset (Fig. 3B) showed that 623 genes supported *Ancistrocladus+Drosophyllum* sister to all other carnivorous plants (hereafter referred to as carnivory supermatrix topology) while 614 genes supported *Ancistrocladus+Drosophyllum* sister to *Nepenthes alata*+*Nepenthes ampullaria* (hereafter referred to as carnivory coalescent topology; Figs. 3A & 4D). Two genes (cluster575 and cluster3300) contributed disproportionately to the overall likelihood.

**Figure 4.**
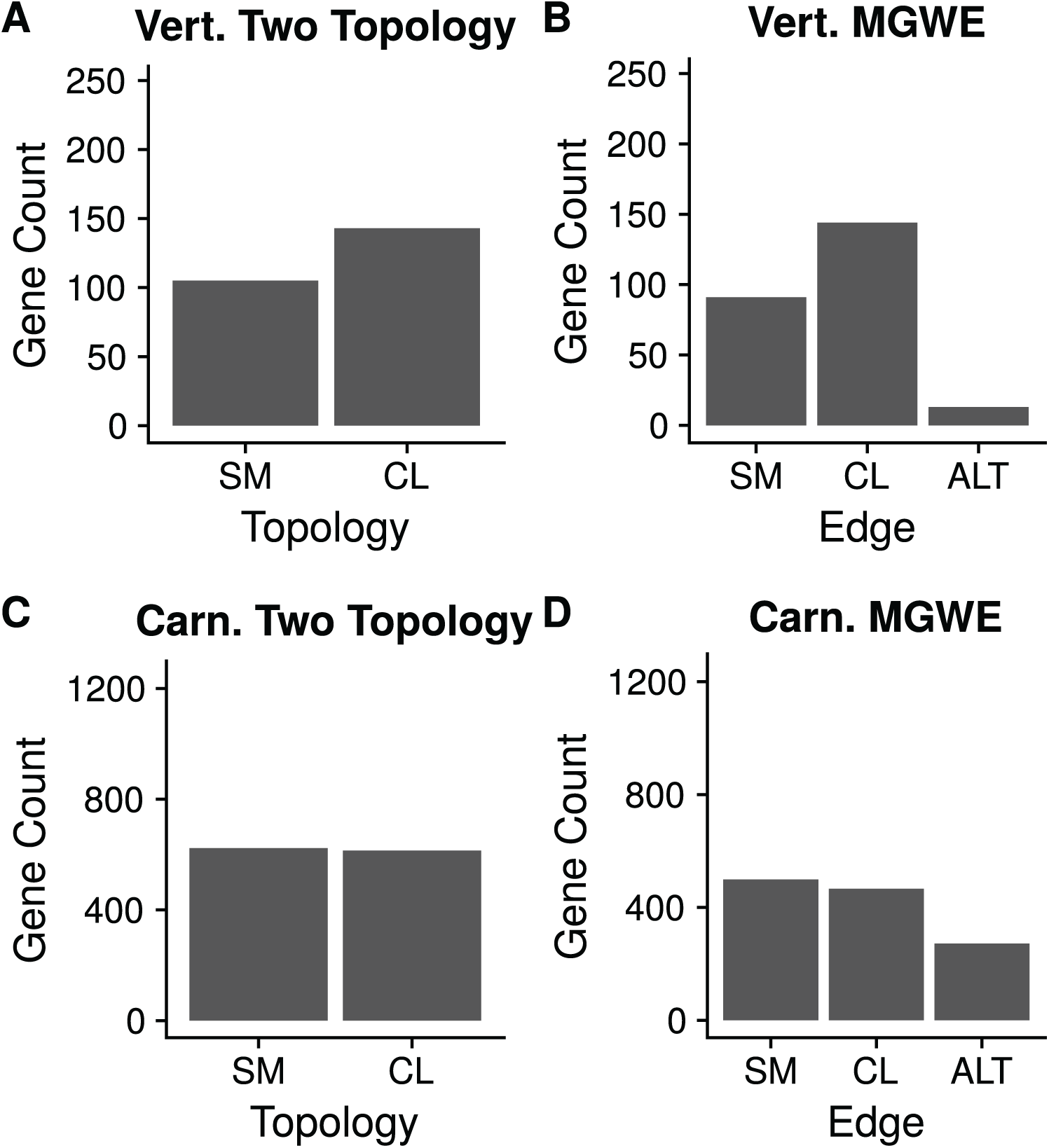
Bar plot representing gene counts for the two-topology and MGWE methods. (A and C) The counts of genes that support the supermatrix inferred maximum likelihood (ML) topology and the coalescent-based maximum quartet support species tree (MQSST), for the vertebrate and carnivory datasets respectively. (B and D) The results of the MGWE analysis for support of the edge found in the ML analysis, the conflicting edge from the MQSST analysis, and the sum of all genes supporting an alternative conflict from an edge in the TREESET.

Individually these two genes have a difference in log-likelihood scores between the two topologies of 33.06 and 16.63, respectively, and support the carnivory supermatrix topology. When we reanalyzed the supermatrix with cluster575 and cluster3300 removed, the carnivory coalescent topology was recovered, with 100% BS support (Supplementary Fig. 1). The difference between the two topologies in log-likelihood of the non-outlier genes ranged from 0.001 to 12.82 with an average of 2.82 for all genes in the analysis.

### Edge-based analysis

We compared MGWE and two topology gene-wise likelihoods involving the contentious bird, crocodilian, and turtle relationships in the vertebrate dataset (Fig. 4B). We found seven unique topologies with the necessary species coverage to conduct the analyses: five gene tree topologies from Chiari et al. (2012) and the two dominant species tree topologies. The set of seven trees included three major conflicting edges for the relationship in question: the two resolutions found in the supermatrix and coalescent trees, and birds sister to crocodilian+mammals+turtles. Ninety-one genes supported the vertebrate supermatrix edge, 144 genes supported the vertebrate coalescent edge, and 13 genes supported the third conflicting edge (Fig. 4B). When comparing the supermatrix analysis with a single set of branch lengths, to that where branches are unlinked, we found lower AICc values for unlinked branches (Table 1). The MGWE AICc scores for the summed likelihoods of the supermatrix (three source trees), the coalescent (three source trees), and the third conflicting edge (one source tree) were highest for the coalescent edge and out of all tested models the coalescent edge was inferred to be the best (Table 1).

**Table 1.** Results of model comparisons.

|  | Relationship | Type | Likelihood | k | AIC | AICc | ΔAICc |
| --- | --- | --- | --- | --- | --- | --- | --- |
| **Vertebrate** | Supermatrix | linked | -1,047,406.05 | 2261 | 2,099,334.11 | 2,099,389.47 | 21,855.08 |
|  |  | unlinked | -1,031,489.81 | 7186 | 2,077,351.63 | 2,077,925.99 | 391.59 |
|  |  | Edge | -1,031,423.67 | 7186 | 2,077,219.34 | 2,077,793.70 | 259.30 |
|  | Coalescent | linked | -1,047,410.07 | 2261 | 2,099,342.15 | 2,099,397.51 | 21,863.11 |
|  |  | unlinked | -1,031,453.35 | 7186 | 2,077,278.71 | 2,077,853.06 | 318.67 |
|  |  | **Edge** | **-1,031,294.01** | **7186** | **2,076,960.04** | **2,077,534.39** | **0** |
|  | Dominant Alternative | Edge | -1,041,062.40 | 7186 | 2,096,496.81 | 2,097,071.16 | 19,536.77 |
| **Carnivory** | Supermatrix | linked | -13,305,055.20 | 11156 | 26,632,422.40 | 26,632,540.58 | 35,228.47 |
|  |  | unlinked | -13,261,947.29 | 39584 | 26,603,062.59 | 26,604,570.70 | 7,258.59 |
|  |  | Edge | -13,258,387.61 | 39584 | 26,595,943.24 | 26,597,451.35 | 139.24 |
|  | Coalescent | linked | -13,305,130.14 | 11156 | 26,632,572.28 | 26,632,690.46 | 35,378.35 |
|  |  | unlinked | -13,262,019.55 | 39584 | 26,603,207.10 | 26,604,715.22 | 7,403.10 |
|  |  | **Edge** | **-13,258,317.99** | **39584** | **26,595,803.99** | **26,597,312.10** | **0** |
|  | Dominant Alternative | Edge | -13,260,106.83 | 39584 | 26,599,381.67 | 26,600,889.78 | 3,577.67 |

\*In the type column, “linked” represents the supermatrix or coalescent topology with a single set of branch lengths, “unlinked” is the supermatrix or coalescent topology with branch lengths varying among genes, and “Edge” is the MGWE analysis. The top AICc score is bolded.

For the carnivory dataset, we found 168 unique tree topologies to include in the tree set. The 168 tree topologies contained 45 conflicting edges for the relationship in question with 3 dominant edges. The MGWE analyses found 499 genes supported the supermatrix edge, 466 genes supported the coalescent edge, and 272 genes supported 15 additional edges (Figs. 2D, 3E). When we further compared the MGWE AICc scores for the supermatrix (44 source trees), the coalescent (56 source trees), and for the third edge (24 source trees) we found the coalescent edge to have the best AICc score out of all tested models (Table 1).

### Outlier gene examination

For the carnivory dataset, we explored the possibility that the strongly conflicting genes cluster575 and cluster3300 reflected methodological error in the assembly pipeline, as is the case for the genes identified by Brown and Thomson (2017) for the vertebrate dataset. However, both the alignment and inferred phylogram for each gene revealed no obvious problems or potential sources of systematic error (sparse alignment, abnormally long branch lengths, etc.). We also explored whether compositional heterogeneity could explain the strongly conflicting results (i.e., that the relationships were not truly conflicting, but instead incorrectly modeled). However, both RY-coding in RAxML and explicit modeling of multiple equilibrium frequencies (2, 3, or 4 composition regimes) across the tree in p4 v1.0 (Foster 2004) failed to overturn the inferred relationships. We further explored the possibility of misidentified orthology. The inferred homolog tree produced from amino acid data, containing the outlier gene from the nucleotide dataset, had no signs of misidentified orthology or gene duplication and loss (i.e., an ortholog within the homolog amino acid tree). We found that with the slower amino acid data the sequences in the nucleotide cluster575 were inferred as a single monophyletic ortholog within a duplicated homolog (Supplementary Fig. 2). The discrepancies that appeared between the amino acid dataset and the CDS dataset were found to be either different in-paralogs/splice sites maintained during the dataset cleaning procedure or short sequences that were not identified as homologs in the coding DNA sequence (CDS) dataset (Supplementary Table 2 and Supplementary Fig. 2).

## DISCUSSION

Biological processes including substitution saturation, hybridization, horizontal gene transfer, and incomplete lineage sorting can contribute to conflicting signal and may explain both conflict and lack of support widely found in phylogenomic datasets (Salichos et al. 2014; Smith et al. 2015; Kobert et al. 2016). To further complicate the challenges facing phylogenomic analyses, high support values, especially from concatenated analyses, can mask significant underlying conflict (Lee and Hugall, 2003; Ryan et al. 2013; Salichos et al. 2014; Smith et al. 2015; Kobert et al. 2016; Pease et al. 2018). We examined two datasets with extensive conflict involving one or several edges for which small changes in analysis approach or dataset composition altered species tree estimates. Both datasets examined here recovered high support for different topologies based on supermatrix or coalescent species tree analyses.

To address the challenges of conflict and support in phylogenomic datasets, several approaches have been outlined in the literature. In addition to identifying gene tree conflict, these approaches have also highlighted outlier genes that dramatically alter supermatrix analyses (Brown and Thomson 2017; Shen et al. 2017). Both datasets contained genes that exhibited outlier behavior with different topologies inferred depending on the inclusion or exclusion of two genes with disproportionate influence on the likelihood (Brown and Thomson 2017; Walker et al. 2017). In the case of the carnivory dataset, the inferred topology changed with the inclusion or exclusion of just 0.0016% of the genes. The outlier genes in a vertebrate dataset were found to be the result of errors in orthology detection (Brown and Thomson 2017). While the genomic resources were not available to fully examine the carnivorous outlier genes (e.g., we do not yet have synteny or information on gene loss), our analyses did not detect any obvious problems with alignment, compositional heterogeneity, or homology. We found one gene, cluster575, to be an ortholog of a gene that experienced a duplication event prior to the divergence of both ingroup and outgroup taxa (Supplementary Fig. 3). While we could not rule out every possible source of error, we also could not identify a source of methodological error, suggesting the possibility that the disproportionate evolutionary information the gene contains to support the conflicting topology is the result of real (albeit unknown) biological processes.

In addition to the discovery of outlier genes, gene tree analyses and topological examinations have been very informative in the exploration of signal for and against conflicting phylogenetic relationships (Castoe et al. 2009; Smith et al. 2011; Shen et al. 2017). While these analyses can be very helpful in dissecting signal, many assume that a single species tree topology that underlies all genes. For several reasons, this may not be an appropriate model (e.g., hybridization, horizontal gene transfer, and other processes). Conflict among gene trees is common and expected from incomplete lineage sorting, hybridization, and other biological processes. For instance, Jarvis et al. (2014) reported that no gene trees from a genomic data set of 48 species of birds matched the inferred species tree. Furthermore, such a result becomes increasingly likely as sampling breadth (both taxa within a clade as well as the age of the clade itself) increases. The results of a shared-topology analysis may be driven by the resolution of a part of the phylogeny other than the area of interest, as shared-topology analyses condition on fully bifurcating trees that necessarily resolve conflict in the entire tree.

To overcome these limitations, we examined edges across a set of empirically supported candidate topologies, as defined by the set of inferred gene trees and the two dominant species tree hypotheses in question. By examining edges, we accommodate for heterogeneity across the rest of the tree, regardless of the process generating that heterogeneity. The vertebrate gene trees contained three alternative edges for the relationship of interest while the carnivory gene trees contained 45 different edges representing 168 different topologies. Both the MGWE analyses and AICc scores of the vertebrate and carnivory datasets suggested a better fit of the coalescent edge than the supermatrix edge (Table 1). Also, in both cases, we found that the AICc score supported the higher parameterized model, as opposed to a single shared topology and branch lengths. While concatenation is commonly performed using a single set of branch lengths, recent work by Neupane et al. (2018) has also suggested that unlinking branches may be preferred. We do not suggest that the highly parameterized model here is the best model in the universe of possible models, only the best of the ones analyzed.

Our results suggest that future studies may benefit from allowing more heterogeneity than is typically involved in a concatenation analysis. This will require careful examination of the complexity involved in large phylogenomic analyses (e.g., missing data; Stamatakis and Alachiotis 2010). The edge based MGWE analyses facilitate rapid and thorough analysis of the support for relationships across each individual gene. By not conditioning on a single topology for all genes, these analyses can better accommodate the existing heterogeneity between genes while still allowing for edge based investigations. The AIC and AICc analyses allow for more explicit comparisons between the disparate models examined here. Future work could expand on these in several ways. For example, the models explored could potentially have significantly reduced parameters by sharing topologies and branch lengths across some compatible gene regions, including potentially scaling branch lengths proportionally (e.g., as is possible with the -spp option in the program iqtree). Nevertheless, the exploratory analyses presented here provide additional evidence that a simple concatenation approach with these large datasets masks important heterogeneity that can be analyzed further to help inform phylogenetic resolution.

The results presented here contribute to a growing body of literature that addresses how phylogenomic analyses should proceed in the presence of highly influential outlier genes, conflicting topologies, and ever expanding datasets (Wickett et al. 2014; Pease et al. 2016; Brown and Thomson 2017; Shen et al. 2017; Yang et al. 2017). For example, some authors have noted, and it is the case here, that supermatrix analyses may be more susceptible to the problem of strong outliers (Shen et al. 2017; Walker et al. 2017). In these studies, the resolutions inferred using a coalescent method were generally favored. When the dominant process generating gene tree conflict is ILS, coalescent methods should perform better. Some coalescent methods that weigh all gene tree equally (e.g., Mirarab and Warnow 2015), may overcome the problem of outlier genes even if incomplete lineage sorting is not the dominant source of conflict simply by eliminating the disproportionate influence of one or two outlying genes. However, with large and broad datasets, it is more likely that processes in addition to ILS have contributed to gene tree conflict and our ability to accurately reconstruct gene trees may be diminished as we move deeper in the tree of life.

While we continue to uncover the patterns and processes that generate conflicting signal within phylogenomic datasets, it is imperative that we continue to explore ways of dissecting the phylogenetic signal within our datasets. By examining the causes of uncertainty and conflict behind recalcitrant nodes, we can present a more measured confidence, or lack thereof, for particular resolutions. For example, while biological processes most certainly have contributed to the conflict within the datasets examined here, other data set assembly issues (e.g., missing data) may also contribute to conflict and low support in these data sets. For example, while the carnivory dataset had extensive data overlap, the vertebrate dataset only had five gene regions that contained sequence data for every species (Supplementary Table 1). Here we present a framework that focuses on analyzing specific conflicting edges with a MGWE analysis that allows for topological heterogeneity outside of the relationships of interest. This approach accommodates the biological realities of heterogeneity among lineages and throughout a phylogeny in order to address specific questions about an edge of interest. While this is just a small contribution to a growing literature on addressing phylogenomic conflict, as we continue to accommodate more heterogeneity within datasets, we should begin to provide more resolution to important nodes in the tree of life.

## ACKNOWLEDGEMENTS

We would like to thank Alexis Stamatakis for advice on RAxML, Caroline Parins-Fukuchi and Nathanael Walker-Hale for critical discussions on the methodology. We would also like to thank Javier Igea, Oscar Vargas, Greg Stull, and Ning Wang for helpful discussion of the manuscript.

---

**Supplementary Figure 1.**
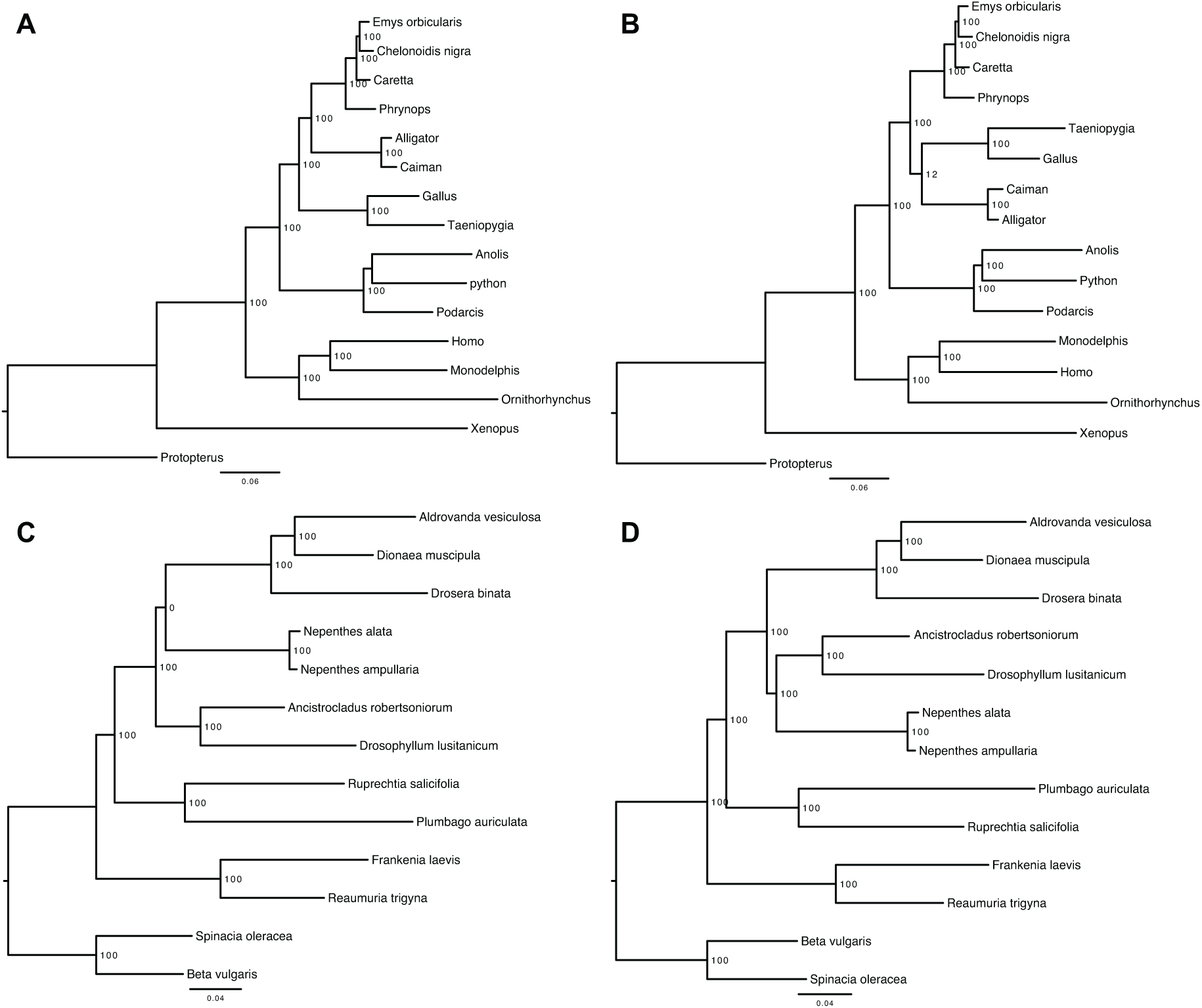
Species trees inferred using maximum likelihood from the different supermatrices. Support at each node was obtained from 200 rapid bootstrap replicates. A) Species tree for vertebrate dataset inferred with all 248 genes included in the supermatrix. B) Species tree for the vertebrate dataset inferred with 8916 and 11434 removed from the supermatrix. C) carnivorous Caryophyllales species tree inferred from all 1237 genes. D) carnivorous Caryophyllales species tree inferred with cluster575 and cluster3300 removed from the supermatrix.

**Supplementary Figure 2.**
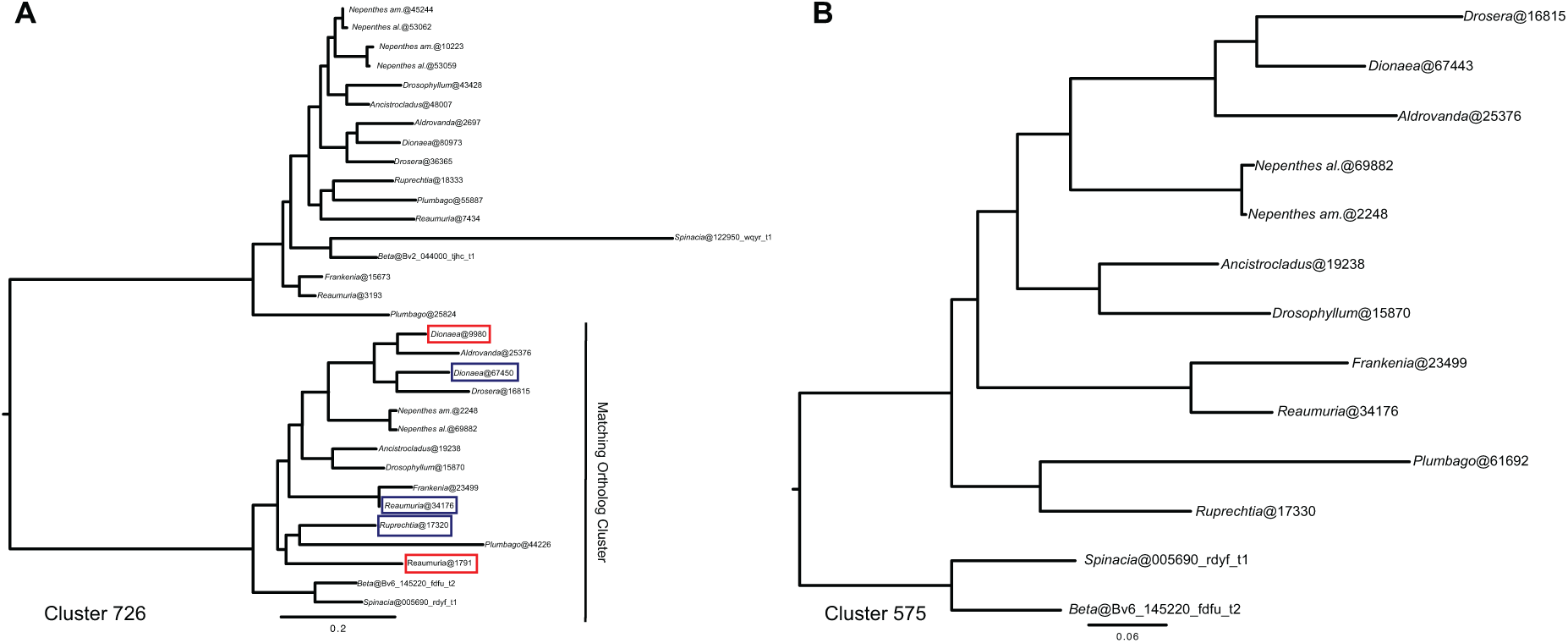
Homolog tree for Amino Acid clustered (726) and CDS clustered (575) highly influential gene in the carnivorous Caryophyllales dataset. Different genes identified in the ortholog clusters are circled on cluster 726. Genes circled in red represent ones that are shorter and were not identified as orthologous in the CDS dataset and genes circled in blue represent alternate paralogs or introsplice sites used between the two clustering analyses.

**Supplementary Table 1.** Number of gene trees in which all the species for a given edges are present. edges correspond to node labels on Fig. 1.

| Edge number | Genes containing all species for the edge |
| --- | --- |
| 0 | 5 |
| 1 | 5 |
| 2 | 246 |
| 3 | 248 |
| 4 | 5 |
| 5 (All turtle, crocodilians, and birds) | 6 |
| 6 | 248 |
| 7 | 6 |
| 8 | 23 |
| 9 | 36 |
| 10 | 45 |
| 11 | 69 |
| 12 | 51 |
| 13 | 94 |
| edge of turtles sister to birds+crocodilians | 36 |

**Supplementary Table 2.** Sources of discrepancy between the orthologs detected in highly influential nucleotide cluster575 and in matching amino acid homolog cluster726.

| Ortholog in 575 | Ortholog in 726 | Seq length of 575 (Nuc) | Seq length of 726 (Nuc) | Reason for misidentification |
| --- | --- | --- | --- | --- |
| Dino@67443<br>(*Dionaea*) | Dino@67450 | 2793 | 2991 | Different copy of the in-paralog or intron splice site was retained |
| Dino@67443<br>(*Dionaea*) | Dino@9980 | 2793 | 510 | Not identified as homologs in blast |
| RuprSFB@17320<br>(*Ruprechtia*) | RuprSFB@17330 | 2787 | 2787 | Different copy of the in-paralog or intron splice site was retained |
| MJM3360@61692<br>(*Plumbago*) | MJM3360@44226 | 2211 | 2403 | Different copy of the in-paralog or intron splice site was retained |
| Retr@34176<br>(*Reaumuria*) | Retr@1791 | 1044 | 546 | Not identified as homologs in blast |

